# Genomic signature of accelerated evolution in a saline-alkaline lake-dwelling Schizothoracine fish

**DOI:** 10.1101/825885

**Authors:** Chao Tong, Miao Li

## Abstract

Tibetan Plateau imposes extremely inhospitable environment on most wildlife. Besides the harsh aquatic environment including hypoxia and chronic cold, high salinity and alkalinity is an increasing threat to Tibetan endemic fishes. Previous genome-wide studies identified key genes contributed to highland fish adaptation to hypoxia and long-term cold, while our understanding of saline and alkaline adaptation in Tibetan fish remains limited. In this study, we performed a comparative genomics analysis in a saline lake-dwelling highland fish *Gymnocypris przewalskii*, aimed to identify candidate genes that contributed to saline and alkaline adaptation. We found elevated genome-wide rate of molecular evolution in *G. przewalskii* relative to lowland teleost fish species. In addition, we found nine genes encoding biological macromolecules associated with ion transport functions underwent accelerated evolution in *G. przewalskii*, which broadly expressed across kidney, gill, liver, spleen, brain and muscle tissues. Moreover, we found putative evidence of ion transport under selection were interacted by co-expression in *G. przewalskii* adaptation to high salinity and alkalinity environment of Lake Qinghai. Taken together, our comparative genomics study identified a set of rapidly evolving ion transport genes and transcriptomic signatures in Schizothoracine fish adaptation to saline and alkaline environment on the Tibetan Plateau.

## 1. Introduction

Environments may shape the genetic landscape of wildlife that inhabit them [1]. The world’s largest and highest highland Tibetan Plateau had undergone continuous uplift during the India-Asia collision since about 45 million years ago, which triggered numerous environmental changes [2,3]. As elevation above sea level increases, a decrease in barometric pressure results in fewer oxygen molecules in the air, which causes hypoxia. Besides, other challenging environments high-altitude dwelling wildlife have encountered are the long-term low temperature and high ultraviolet radiation [4,5]. Understanding how organism adapt to their dwelling environment is central to answering many ecological and evolutionary questions, but it remains a formidable task to fully uncover the mechanisms of adaptive process [6]. Adaptation at the molecular level can occur by adaptive mutation in key genes over prolonged evolutionary time scales [7]. Recent genome-wide studies have identified key genes associated with hypoxia response and energy metabolism in Tibetan terrestrial animals adaptation to high altitude [8–10]. Nevertheless, the adaptive mechanism of Tibetan aquatic organisms to highland water environment is yet well-studied [11].

Unlike Tibetan terrestrial animal, the draft genomes of very few Tibetan aquatic organisms had been sequenced [12,13], the genomic basis of aquatic animals adaptation to water environments at high altitude still remain largely unknown. The Schizothoracine fishes are the predominant aquatic fauna on the Tibetan Plateau, which had evolved specific phenotypic characteristics to adapt to the harsh aquatic environments, such as hypoxia, chronic cold and high salinity and alkalinity. Comparative genomics approaches have the power to facilitate investigation of the genomic basis of evolution and adaptation [14]. Recent comparative genomics studies based on transcriptomic data of several Schizothoracine species have identified a number of candidate genes that underwent positive selection during the long-term adaptive processes to harsh environments on the Tibetan Plateau, such as BYSL and HSF1 associated with hypoxia response [15] and ND1, ATAD2 and ARL3 that involved into cold response [16,17]. Notably, an increasing number of lakes are existing or towards saline and alkaline due to the geological evolution and global climate changes on the Tibetan Plateau [3,18]. For instance, Lake Qinghai, the largest salt lake in China, is highly saline (up to 13‰) and alkaline (up to pH 9.4) water environment. It is also a typical salt lake with unusually high sodium, potassium and magnesium concentration [18,19]. Intriguingly, Lake Qinghai used to be freshwater and connected to the Yellow River, while was late separated with the upper reaches of the Yellow River during the geological event “Gonghe Movement” (approximately 15 mya) [19,20]. Moreover, the increasing of water salinization is a growing threat to freshwater fish species [21,22]. Tibetan freshwater endemic fishes are long suffering these harsh conditions challenges [11,18]. The main focus of the genetic mechanism of highland adaptation in Tibetan fish are still on hypoxia and chronic cold response [15,23–25]. However, the genomic signature of high salinity and alkalinity adaptation in Schizothoracine fish have yet to be comprehensively determined.

Unlike other broadly distributed Schizothoracinae fish species, *Gymnocypris przewalskii* is only endemic to Lake Qinghai [19,20,26]. Past studies suggested that *G. przewalskii* has gradually evolved from freshwater fish to tolerate high salinity and alkalinity of Lake Qinghai during the early to late Holocene [26]. Because of the unique evolutionary history in Lake Qinghai at high altitude, *G. przewalskii* provides an exceptional model to investigate the genetic mechanisms underlying adaptation to high salinity and alkalinity environment on the Tibetan Plateau. In this study, we performed a comparative genomics analysis and identified a set of ion transport genes that showing strong signals of rapidly evolving in *G. przewalskii*. Specifically, we used the *de novo* transcriptome assemblies from multiple tissue RNA-seq data and five well-annotated teleost fish genomes for comparison. In addition, we estimated the genome-wide nucleotide substitution rate of each fish species. Moreover, using the tissue-transcriptomics, we characterized the expression patterns of rapidly evolving ion transport genes in kidney, gill, liver, spleen, brain and muscle of highland fish, *G. przewalskii*.

## 2. Materials and methods

### 2.1. Data collection and transcriptome assembly

We downloaded the transcriptome sequencing data of Schizothoracine fish *G. przewalskii* from NCBI SRA database (https://www.ncbi.nlm.nih.gov/sra). Specifically, we collected six tissues transcriptomics including kidney, gill, liver, spleen, brain and muscle of *G. przewalskii* (supplementary table S1). At first, we checked the quality of the raw sequencing reads using FastQC v 0.11.8 (https://www.bioinformatics.babraham.ac.uk/projects/fastqc/). Sequencing adapters and reads with a quality score < 20 were trimmed with Trimmomatic v.0.36 [27], resulting in clean sequencing reads. Then, we performed *de novo* transcriptome assembly using Trinity v2.8.5 [28] with default parameters. After assembly, the redundant transcripts were removed using CD-HIT v4.8.1 [29] with the threshold of 0.90, and only the longest transcript under each cluster was extracted as unigene (unique gene). Next, we predicted the open reading frame (ORF) of each unigene using TransDecoder (https://github.com/TransDecoder/TransDecoder). Finally, we translated the nucleotide sequences of protein-coding genes from the assemblies of *G. przewalskii* into amino acid sequences using an in-house-developed perl script.

### 2.2. Orthologs identification and sequence alignment

We included five well-annotated teleost fish genomes for comparative genomics analysis and downloaded from Ensembl database (http://useast.ensembl.org/index.html), including zebrafish (*Danio rerio*), tilapia (*Oreochromis niloticus*), medaka (*Oryzias latipes*), fugu (*Takifugu rubripes*) and cod (*Gadus morhua*). Then, we built a local protein database including the sequences from above five fish genomes and *G. przewalskii* transcriptome assemblies. Next, we downloaded the curated orthology map of Actinopterygii (ray-finned fish) from OrthoDB database (release 8) [30] which contains 21,952 orthologous gene groups information. Of these seed orthologous groups in HaMStR v13.2.6 [31], we identified the orthologs in each fish species with E-values of less than 10^−20^. Moreover, we aligned and trimmed the protein sequences of the orthologous groups using PRANK [32] and MATFF v 7.450 [33], and trimmed using trimAl v1.2 [34] with the parameter “-automated1”. Among the identified orthologs, we identified one-to-one, one-to-many and many-to-many orthologs in each fish species. For each 1:1 orthologous pair (i.e. genes for which only one gene from each species matches the given OrthoDB orthologous gene group), we only selected the longest transcript associated with the gene for each pair of species as putative single-copy ortholog. Finally, we identified the core single-copy orthologs that were shared by above six fish species.

### 2.3. Genome-scale concatenation and coalescent based dataset construction

We performed the alignment of core shared single-copy orthologs of six fish species using MUSCLE [35] with default parameters and trimmed using trimAl [34] with parameter “-automated1”. In addition, we filtered the core shared single-copy orthologs with strict constraints, including length (minimum 200 aa) and sequence alignment (maximum missing data 50% in alignments). Next, we prepared two types of datasets after filtration. Firstly, we concatenated all core shared single-copy genes from each species into one-line sequence as a supergene using a custom python script (genome-scale concatenation-based dataset), respectively. Secondly, we conducted a genome-scale coalescent-based dataset including core shared single-copy genes from each species.

### 2.4. Molecular evolution analysis

We used the clipped species tree (Figure 1A) including above six fish species from a previous study [16]. To estimate lineage-specific evolutionary rates for each fish species, we aligned core shared single-copy orthologs using MUSCLE [35], derived nucleotide alignments from protein alignments using PAL2NAL v14 [36], and estimated pairwise non-synonymous to synonymous substitutions (dN/dS) of nucleotide alignments using the CodeML package in PAML 4.7a [37]. Specifically, we used the free-ratio model (“several ω ratio”) to calculate the ratio of dN to dS nucleotide changes separately for each ortholog and a concatenation of all alignments of single-copy orthologs from above six fish species. Parameters, including dN, dS, dN/dS, N*dN, and S*dS values, were estimated for each branch, and genes were discarded if N*dN or S*dS < 1, or dS > 2, following previous studies [11,16,17].

**Figure 1.**
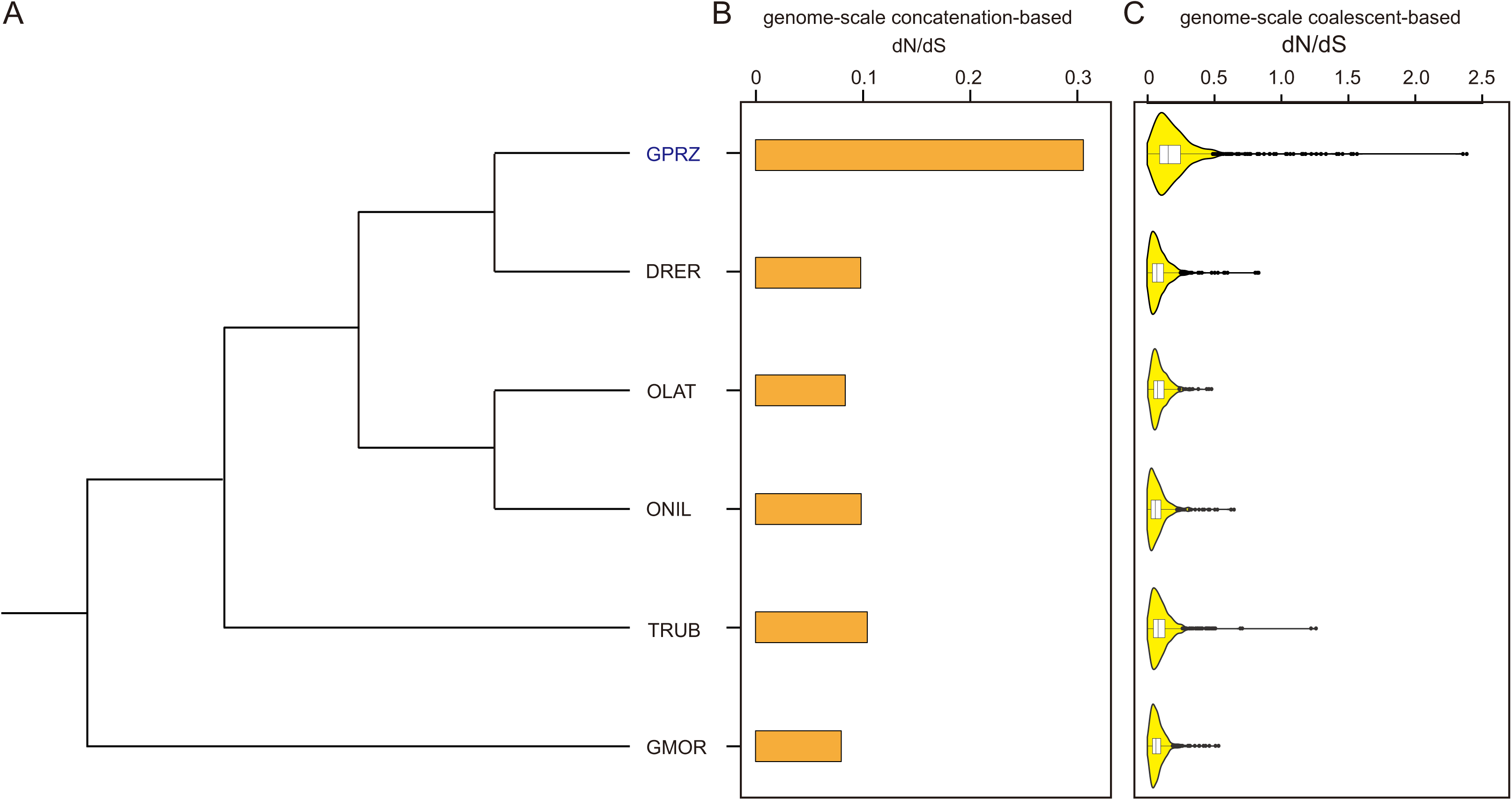
Genome-wide molecular evolutionary feature of six fish species. (A) Clipped species tree used in this study. GPRZ, *Gymnocypris przewalskii*, DRER, *Danio rerio*, OLAT, Oryzias latipes, ONIL, Oreochromis niloticus, TRUB, Takifugu rubripes, GMOR, *Gadus morhua*. (B) Barplot depicting the dN/dS of concatenated supergenes in each fish species. (C) Violin plot depicting the dN/dS of each coalescent orthologs in each species estimated by free-ratio model.

We sought to identify a set of genes with elevated dN/dS in *G. przewalskii* relative to other five fish species. At first, we ran two branch models using CodeML package in PAML 4.7a [37] to identify rapidly evolving genes (REGs) in *G. przewalskii* lineage with corresponding nucleotide alignments, specifically with the null model assuming that all branches have been evolving at the same rate and the alternative model allowing the focal foreground branch (*G. przewalskii*) to evolve under a different evolutionary rate. Next, we used a likelihood ratio test (LRT) in R software, package MASS with df□=□1 to discriminate between the alternative model and the null model for each single-copy orthologs in the genesets. We only considered the genes as rapidly evolving with a significantly faster rate in the foreground branch if the adjusted *P* value□<□0.05 and higher dN/dS in the focal foreground branch than focal background branches (other four fish species). Finally, we annotated the rapidly evolving genes with gene ontology (GO) function category using R software, package topGO [38].

## 2.5. Gene expression analysis

After preparation of clean reads from six tissue-transcriptomics (kidney, gill, liver, spleen, brain and muscle) of *G. przewalskii*, we performed the transcript quantification by mapping all clean reads to the transcriptome assemblies using RSEM v1.3.1 [39] to obtain expected counts and fragments per kilobase million (FPKM). In addition, we primarily focused on the expression pattern of rapidly evolving genes (REGs), and calculated the FPKM value of each REG in each tissue. At last, we annotated the differentially expressed REGs by gene ontology using R software, package TopGO [38].

## 3. Results

### 3.1. Transcriptome assemblies and orthologs

By pooling six tissue-transcriptome sequencing data, the de novo transcriptome assembly of *G. przewalskii* yielded 409,685 transcripts, with an N50 of 1,796 bp and an average length of 986 bp. After removing redundant isoforms and extraction of longest isoform among alternative transcripts, a total of 357,601 unigenes were obtained, with an N50 of 3,079 bp and a mean length of 1,992 bp. After protein-coding gene prediction with TransDecoder, we totally obtained 137,539 unigenes with full or partial length of gene coding regions (CDS) in *G. przewalskii*.

After identification of orthologs according to the curated orthologous gene groups of Actinopterygii in each fish species, we obtained a total of 74,107 putative orthologs in 16,379 orthologous gene groups (Table 1). After strict 1:1 ortholog selection, we identificated 16,379 longest orthologs that represent their gene groups as unique ortholog (Table 1). In addition, we eventually obtained core 10,260 orthologs that shared by all six fish species, making them suitable for comparative genomics analysis.

**Table 1.** Summary of orthologous genes in five fish genomes and *G. przewalskii* transcriptomic assemblies.

| Species | Genes | Genes in orthologous groups | Unique orthologs |
| --- | --- | --- | --- |
| *Danio rerio* | 52,089 | 29,232 | 17,001 |
| *Gadus morhua* | 22,100 | 16,884 | 16,390 |
| *Takifugu rubripes* | 47,841 | 25,137 | 16,071 |
| *Oryzias latipes* | 24,674 | 17,857 | 15,877 |
| *Oreochromis niloticus* | 26,763 | 19,432 | 17,433 |
| *Gymnocypris przewalskii* | 137,539 | 74,107 | 16,379 |

### 3.2. Genome-wide nucleotide substitution rate

After estimation of the nucleotide substitution rates of each branch that represented each fish species based on 6,742 core shared single-copy orthologs, we found that Schizothoracine fish *G. przewalskii* had elevated terminal genome-wide concatenation-based dN/dS compared to other five fish species (Figure 1B). Furthermore, we also found similar elevated pattern of genome-wide coalescent-based dN/dS in *G. przewalskii* relative to other species (Figure 1C).

### 3.3. Rapidly evolving genes

A set of genes with the signature of an increase rate of non-synonymous changes and underwent accelerated evolution, namely rapidly evolving genes. We identified 466 putative rapidly evolving single-copy orthologs (REGs) in *G. przewalskii* (supplementary table S2). Among this set of genes, the most interesting finding was REGs included genes associated with ion transport functions. This group included sodium channel subunit beta-3 (SCN3B), solute carrier family 13 member 3 (SLC13A3), transmembrane protein 175 (TMEM175) and H(^+^)/Cl(^−^) exchange transporter 7 (CLCN7) (Table 2). Moreover, we found a number of REGs associated with mitochondrial function and also involved ion transport process, such as sodium/potassium-transporting ATPase subunit beta-2 (ATP1B2), calcium uniporter protein (MCU) and calcium uptake protein 2 (MICU2) (Table 2). Besides the ion transport genes, we found a large number of genes involved energy metabolism function, such as ATP5c1 and ATP5b associated with ATP binding and oxidative phosphorylation process (supplementary table S2). Although previous comparative genomics studies with highland fish identified several candidate genes with the signals of positive selection [15,23,24], here, we failed to identify any gene that potentially associated with hypoxia response.

**Table 2.** List of rapidly evolving ion transport genes in *Gymnocypris przewalskii*.

| Gene name | Description | Adjusted P-value |
| --- | --- | --- |
| SCN3B | Sodium channel subunit beta-3 | 0.000076 |
| ATP1B2 | Sodium/potassium-transporting ATPase subunit beta-2 | 0.020563 |
| NALCN | Sodium leak channel non-selective protein | 0.029246 |
| SLC13A3 | Solute carrier family 13 member 3 | 0.002234 |
| TMEM175 | Transmembrane protein 175 | 0.002100 |
| CLCN7 | H(+)/Cl(-) exchange transporter 7 | 0.000019 |
| TRPV6 | Transient receptor potential cation channel subfamily V member 6 | 0.022243 |
| MCU | Calcium uniporter protein, mitochondrial | 0.003946 |
| MICU2 | Calcium uptake protein 2, mitochondrial | 0.002823 |

### 3.4. Tissue expression patterns of rapidly evolving ion transport genes

After mapping the clean reads from six tissue-transcriptome sequencing data, we obtained the expression level of each unigenes based on FPKM value (supplementary table S3). We focused on the expression pattern of ion transport genes with the signature of accelerated evolution. Notably, we found eight rapidly evolving ion transport genes were broadly expressed in six tissues, except *transient receptor potential cation channel subfamily V member 6* (*TRPV6*) that only expressed in liver (Figure 2A). In addition, the hierarchical clustering which illustrated by heatmap indicated that four genes (*ATP1B2*, *MCU*, *CLCN7* and *NALCN*) and another five genes (*MICU2*, *SCN3B*, *TMEM175*, *TRPV6* and *SLC13A3*) showed similar tissues expression patterns, respectively (Figure 2B). Moreover, gene ontology (GO) enrichment analysis showed that this set of differentially expressed REGs was significantly enriched multiple functions, such as ion transport (GO:0006811, *P* = 0.00031), sodium ion transport (GO:0006814, *P* = 0.00047), calcium ion transport (GO:0006816, *P* = 0.00056), chloride transport (GO:0006821, *P* = 0.00067), response to pH (GO:0009268, *P* = 0.00069) and response to calcium ion (GO:0051592, *P* = 0.00078) (Figure 2C).

**Figure 2.**
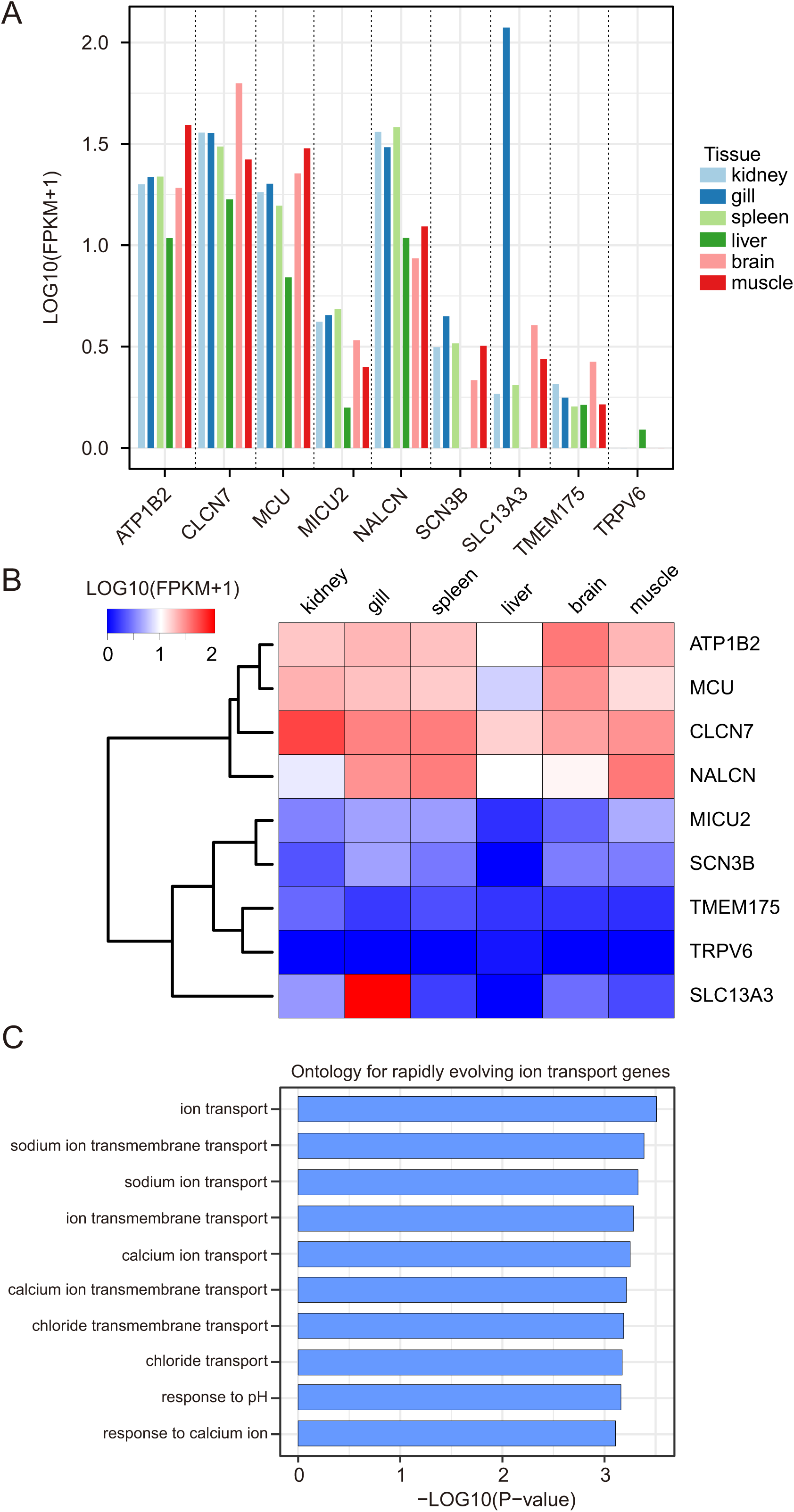
Expression feature of rapidly evolving ion transport genes (REITGs) in six tissues of *G. przewalskii*. (A) Barplot depicting the expression level of nine REITGs in kidney, gill, liver, spleen, brain and muscle tissues based on Log_10_(FPKM + 1) value which estimated from RNA-seq data. (B) Heatmap depicting the expression level comparison of each REITG based on Log_10_(FPKM + 1) values. Tissue type and gene name are shown on the y-axis and x-axis, respectively. Plot colors reflect the expression level, ranging from low (blue) to high (red). (C) Barplot depicting the significantly enriched gene ontology for differentially expressed REITGs.

## 4. Discussion

Over the past few years, transcriptome-based assembly approach enables comparative genomics studies widely employed in many Tibetan endemic organisms to provide insights of highland adaptation [15–17,23,40,41]. Unlike whole genome data, although transcriptome sequencing is an effective and accessible approach to initiate comparative genomic analyses on non-model organisms [28], it still can not cover the whole protein coding gene repertoire of one species. Previous transcriptome studies on Tibetan fishes mainly included one or two tissues [17,40,41], our present study included six tissues (kidney, gill, liver, spleen, brain and muscle) RNA-seq data of *G. przewalskii* and generated much more transcripts than previously assemblies [16,40]. In addition, we used curated orthology mapping approach [42] and identified more than 15,000 pairwise orthologous genes in each fish species and over 10,000 core single-copy orthologs shared by six species, which included much more orthologs than our early studies [16,17]. These putative single-copy orthologs are the important bases for comparative genomic analysis. Notably, most high-altitude dwelling Schizothoracine fish species are polyploidy with high complexity and large size of genomes, the whole genome data is long lacking [11]. Therefore, comparative genomic analysis based on transcriptome assemblies of Schizothoracine fish will still be the tendency in recent years.

Our present study pinpointed that highland fish, *G. przewalskii* has elevated rate of molecular evolution (dN/dS) on both concatenation and coalescent genomic-scales compared with lowland fish species, indicating that *G. przewalskii* may be under rapidly evolving. Not surprisingly, this result was consistent with previous studies in other Tibetan endemic fish species [15–17,23,41]. In addition, this finding highlighted animals endemic to the Tibetan Plateau underwent accelerated evolution (high dN/dS) relative to low-altitude dwelling organisms [9,10]. Furthermore, species inhabit similar ecological niches may be shaped by convergent evolution to form physiological or morphological similarities [43]. Like other Tibetan terrestrial wildlife, our finding implied that the elevation of genome-wide nucleotide substitution rate is one of adaptive process of *G. przewalskii* to harsh environment in Lake Qinghai, including the increasing of water salinization.

Accelerated evolution at molecular level may be reflected by an increased rate of non-synonymous changes within genes involved in adaptation [44]. Our present comparative study identifies a set of rapidly evolving genes associated with ion transport function in *G. przewalskii*. These genes encoded biological macromolecules which mainly functioning in sodium ion transport, calcuim ion transport, chloride transport and response to pH processes. This result is consistent with findings in an extremely alkaline environment dwelling fish, *Leuciscus waleckii* [45], indicating that the alkaline environment of both Lake Qinghai and Lake Dali Nur spurred accelerated evolution of ion transport genes in both fish species. Notably, the rapidly evolving gene repertoire of *G. przewalskii* included SLC13A3, TMEM175 and CLCN7. Solute carrier (SLC) is a family that encoded transmembrane transporters for inorganic ions, amino acids, neurotransmitters, sugars, purines and fatty acids, and other solute substrates [46]. Past evidence suggested that the adaptive evolution of solute carrier genes contribute to high salinity and alkalinity adaptation in fishes [45–47]. Specifically, SLC13 is a subfamily of sodium sulphate/carboxylate cotransporters [48]. Moreover, CLC gene is a family of H^+^/Cl^−^ exchange transporter that mediate transmembrane Cl− transport [49]. In addition, previous study suggested that TMEM175 is involved in potassium channel activity [50]. Therefore, we suggested that ion transport genes underwent rapidly evolving is another adaptation strategy for *G. przewalskii* to cope with the severe saline and alkaline stress in Lake Qinghai.

Previous studies identified a number of genes under accelerated evolution that mainly involved energy metabolism pathways [11,15–17,23–25]. Compared with a few rapidly evolving ion transport genes that were found in *G. przewalskii*, this present study identified a number of candidate genes that related to energy metabolism and contributed to long-term cold adaptation. Gene associated with energy metabolism showing signs of adaptive evolution is one of the genomic signatures that had been identified in Tibetan Plateau dwelling animals [16,17]. Our finding is consistent with previous comparative genomics studies in highland fishes as well [15–17]. A set of genes functioning in energy supply and metabolism were under accelerated evolution in *G. przewalskii*, such as ATP5b and ATP5c, ATP synthase subunit beta. In addition, although hypoxia adaptation is one of the significant adaptive processes contributed to highland adaptation in endemic animals that dwelt at high altitude [10,16,17], we still were not able to identify any rapidly evolving genes associated with hypoxia response function in present study. Indeed, there is a long controversial issue about hypoxic environment and hypoxia response for Tibetan fish species. Obviously, more physiological, ecological and genomic analysis were required to reveal the mechanism of highland fish adaptation to hypoxia.

A set of previous studies indicated that fish gills, kidney, liver and spleen are key tissues that contributed to saline and alkaline tolerance [51,52]. Using tissue-transcriptomic data, we characterized the expression profiles of six tissue types. Most of rapidly evolving ion transport genes have broad expression patterns across all tissues. In addition, these broadly expressed ion transport genes were mainly associated with sodium ion transport, chloride transport and response to pH function by gene ontology annotation. This finding indicates that ion transport genes in *G. przewalskii* experiencing accelerated evolution may have general functions and involve into multiple biological processes. Furthermore, we found a set of rapidly evolving ion transport genes that involved distinct pathways showed the similar tissue expression patterns. That is said, these ion transport genes under selection were putatively interacted by cooperation in *G. przewalskii* adaptation to high salinity and alkalinity environment of Lake Qinghai. Therefore, this finding indicated that future Schizothoracine fish comparative genomics study, including increasing sequencing and function assay, can further clarity the molecular basis of saline and alkaline adaptation of high-altitude dwelling fishes.

## 5. Conclusion

We used comparative genomics based on the *de novo* assemblies from pooled six tissues transcriptomes to identify the genomic signature of saline and alkaline adaptation in a highland fish, *G. przewalskii*. These putative genomic signatures included: (1) Schizothoracine fish, *G. przewalskii* had an elevated genome-wide nucleotide substitution rate than lowland teleost fish species; (2) a number of genes associated with ion transport and energy metabolism functions were found in *G. przewalskii* with elevated molecular evolutionary rate (dN/dS) showing the signature of rapidly evolving; (3) most of rapidly evolving ion transport genes associated with sodium ion transport, calcium ion transport and chloride transport were broadly expressed in kidney, gill, liver, spleen, brain and muscle of *G. przewalskii*; (4) A set of rapidly evolving ion transport genes exhibited similar tissue expression patterns and were interacted by co-expression in *G. przewalskii*. Altogether, our present study will provide the genomic signatures of rapidly evolving ion transport genes, and gain the insights into the saline and alkaline adaptation of high-altitude dwelling fishes.

## Acknowledgments

This work was funded by the Field Research Fund of University of Pennsylvania Biology Department.

